# Train-the-Trainer as an Effective Approach to Building Global Networks of Experts in Genomic Surveillance of AMR

**DOI:** 10.1101/2021.06.18.448940

**Authors:** Monica Abrudan, Alice Matimba, Dusanka Nikolic, Darren Hughes, Silvia Argimón, Mihir Kekre, Anthony Underwood, David Aanensen, The NIHR Global Health Research Unit (GHRU) on Genomic Surveillance of Antimicrobial Resistance

## Abstract

Advanced genomics and sequencing technologies are increasingly becoming critical for global health applications such as pathogen and antimicrobial resistance (AMR) surveillance. Limited resources challenge capacity development in low- and middle-income countries (LMICs), with few countries having genomics facilities and adequately trained staff. Training research and public health experts who are directly involved in the establishment of such facilities offers an effective, but limited, solution to a growing need. Instead, training them to impart their knowledge and skills to others provides a sustainable model for scaling up the much needed capacity and capability for genomic sequencing and analysis locally with global impact. We designed and developed a Train-the-Trainer course integrating pedagogical aspects with genomic and bioinformatics activities. The course was delivered to 18 participants from 12 countries in Africa, Asia, and Latin America. A combination of teaching strategies culminating in a group project created a foundation for continued development at home institutions. Upon follow-up after 6 months, at least 40% of trainees had initiated training programs and collaborations to build capacity at local, national, and regional level. This work provides a framework for implementing a training and capacity building program for the application of genomics tools and resources in AMR surveillance.

**40-word summary:** This work provides a framework for implementing a training and capacity building program for the application of genomics tools and resources in AMR surveillance. We outline a Train-the-Trainer course integrating pedagogical aspects with genomic and bioinformatics activities.

## INTRODUCTION

### Capacity Building for Genomic Surveillance of AMR

Antimicrobial Resistance (AMR) is a major global challenge that is increasing the burden on health systems and impacting on food sustainability, environmental wellbeing, and socio-economic development globally [1]. To strengthen knowledge and evidence base and inform decision-making, the World Health Organization (WHO) have recommended the application of advanced technologies such as whole genome sequencing (WGS) for surveillance of pathogens and AMR. However, in low- and middle-income countries (LMICs), implementation of genomic surveillance of AMR is challenged by the high costs of infrastructure and equipment, and the lack of specialized workforce [2].

To build capacity for pathogen surveillance and genomics, a critical mass of expertise in sequencing techniques and bioinformatics is essential. The establishment of local and regional networks and trainers will raise the impact and ensure the sustainability of the long-term implementation of the genomic surveillance of AMR.

In 2018, the Centre of Genomic Pathogen Surveillance (CGPS) was funded as a partner within the NIHR Global Health Research Unit (GHRU) on Genomic Surveillance of Antimicrobial Resistance, with the overall aim to establish intelligent genomic surveillance of bacterial pathogens through appropriate sampling and analysis with partners in Colombia, India, Nigeria, and the Philippines. To address the need for expertise, in 2019, the GHRU collaborated with Wellcome Genome Campus Advanced Courses and Scientific Conferences (ACSC) to design and deliver a Train-the-Trainer (TtT) course, which focused on the application of genomics in AMR surveillance [3]. The goal was to establish a global network of trainers, capable of delivering direct training and presentation of concepts to varied audiences and contributing to building capacity for the genomic surveillance of AMR laboratories in LMICs.

### An Effective Method for Building Sustainable Global Networks of Experts

Train-the-Trainer (TtT) refers to a program or course where subject domain specialists receive training in a given subject and pedagogical skills needed to train and share their expertise with others. Scaling up training programs in various domains has improved coverage and access, reduced costs, utilized local trainers who are knowledgeable about local issues, and encouraged collaboration among researchers and staff from neighboring research facilities [4].

Several organizations have developed networks of training expertise aimed at addressing the knowledge gaps in fast-changing fields, such as genomics and bioinformatics. TtT courses teach researchers skills to train others in topics that are broad in scope, such as data analysis (e.g. Carpentries), bioinformatics (e.g. H3ABioNet in Africa, EBI-CABANA in Latin America, Next Generation Sequencing Bioinformatics program in Australia) and digital skills (ARDC, a national initiative to support Australian research) [5-9].

The ACSC Program also developed tailored models of training trainers, which were delivered in addition to the usual training in biomedicine and genomics for researchers and healthcare professionals worldwide. Participants were equipped with tools for training others following good educational practice. A key outcome of the ACSC TtT program was the establishment of regional instructor teams, who collaborate with international experts to deliver training in LMIC regions where the need for trainer expertise is most critical.

### Course Development

The goal of the TtT course was to equip participants with skills for the application of effective methods for training adult learners in laboratory techniques and bioinformatics required for the implementation of genomic surveillance of AMR towards establishing global networks of expertise. To effectively train others, participants’ learning outcomes included demonstration of knowledge in: a) setting up a genomic surveillance laboratory and bioinformatics pipelines; b) implementation of laboratory and bioinformatics procedures for generation, analysis, and interpretation of genome data for AMR surveillance; and c) teaching genomic surveillance of AMR to varied audiences.

The course instructor team included experts in education, bioinformatics, epidemiology, genomics, and AMR laboratory methods. Prior to developing the course, the instructor team undertook a TtT course, which provided guidance for the development of an integrated program encompassing pedagogical aspects, and laboratory and computational techniques.

The course was attended by 18 scientists, working in academic research centers or national reference laboratories in 12 countries (Argentina, Brazil, Colombia, Cuba, India, Malaysia, Nigeria, South Africa, Sudan, Thailand, the Philippines, and Vietnam).

### The Structure of the Course

The course consisted of 12 modules, which were run either in parallel or as joint sessions (Figure 1). The course ran over 6 days and included didactic lectures, interactive sessions, and seminars, totaling 47.5 contact hours.

**Figure 1.**
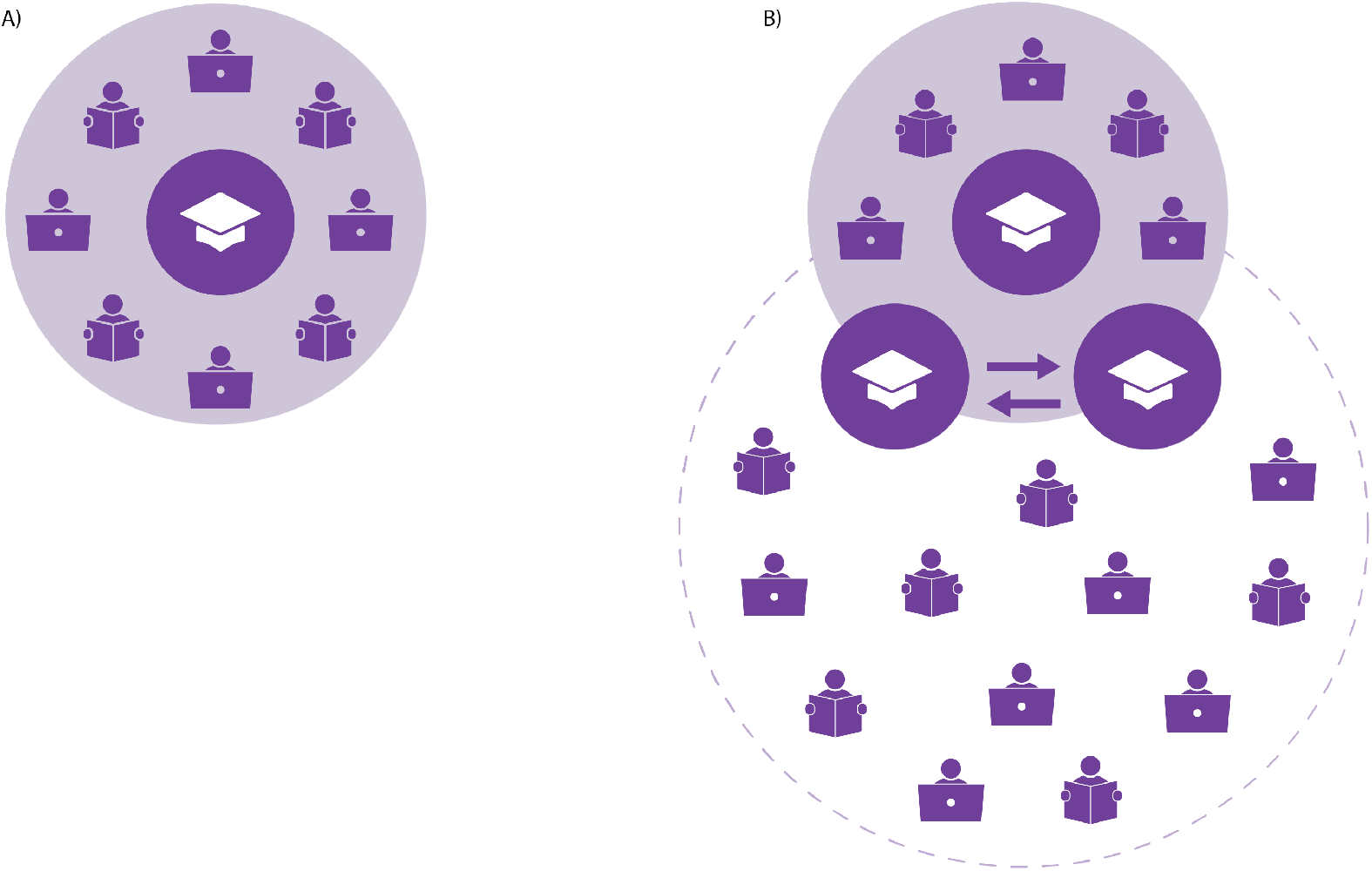
Figure shows the difference in sizes of key beneficiaries of training, between A) conventional training courses, where a trainer (mortarboard hat) teaches participants that gain individual skills, which they then apply to their own data (the audience is shown as people with books and laptops); and B) Train-the-Trainer courses, where the trainer (mortarboard hat) teaches other trainers, who go on to teach more people (audience), and the final audience is larger.

The pedagogical modules and refresher sessions were run with the whole cohort of attendees. Later, the participants were split into two workstreams specific to their interests and expertise. The Laboratory workstream covered processes that enable WGS of bacterial pathogens, from sample collection to sequencing quality assurance. The Bioinformatics workstream covered various aspects, from building the infrastructure and pipelines required for WGS data analysis, to interpretation applied to global contexts. The course concluded with a session in which participants learned aspects needed to be considered when communicating with policymakers and other decision makers. Four guest lectures given by scientists and public health experts in AMR surveillance offered complementary views on the challenges faced in the implementation of genomic AMR surveillance in the UK versus LMICs.

Participants were introduced to a Training and Learning Toolkit, an online book of worksheets, where they collected their own work during the course, as well as their notes on course/module design, networks, resources, and reflections.

### A Strong Emphasis on Pedagogy

The TtT course was designed to support social learning and teaching, where both the instructors that facilitated the training and course participants took active roles and responsibilities for their own teaching and learning. Learning and teaching took several formats: direct teaching/training through presentation or demonstration followed by Q&A session (for example, direct instruction on use of scripts/programs, on setting up labs, or on different lab procedures); group work (for peer-learning on any topic covered in the course); individual reflection, and consolidation. The course considered training on many levels, and the course instructors and facilitators modelled behavior to be adapted and adopted by the participants in the last module of the course, but also in their future teaching and learning.

The expanded learning outcomes (LOs) and module content purposefully integrated the various theoretical and practical aspects of both the pedagogy and the scientific training (Table 1). Learning outcomes 1-4 were covered in the pedagogy sessions introducing educational theory and active learning pedagogy. Topics included elements of course design, writing LOs, and assessments. Scientific content focused on LOs 5 and 6, providing refresher content and demonstration of key techniques and pipelines required for effective AMR genomic surveillance. This further developed to higher-level outcomes 7 and 8, where participants collaboratively designed and presented their modules combined with feedback from the instructors.

**Table 1.**
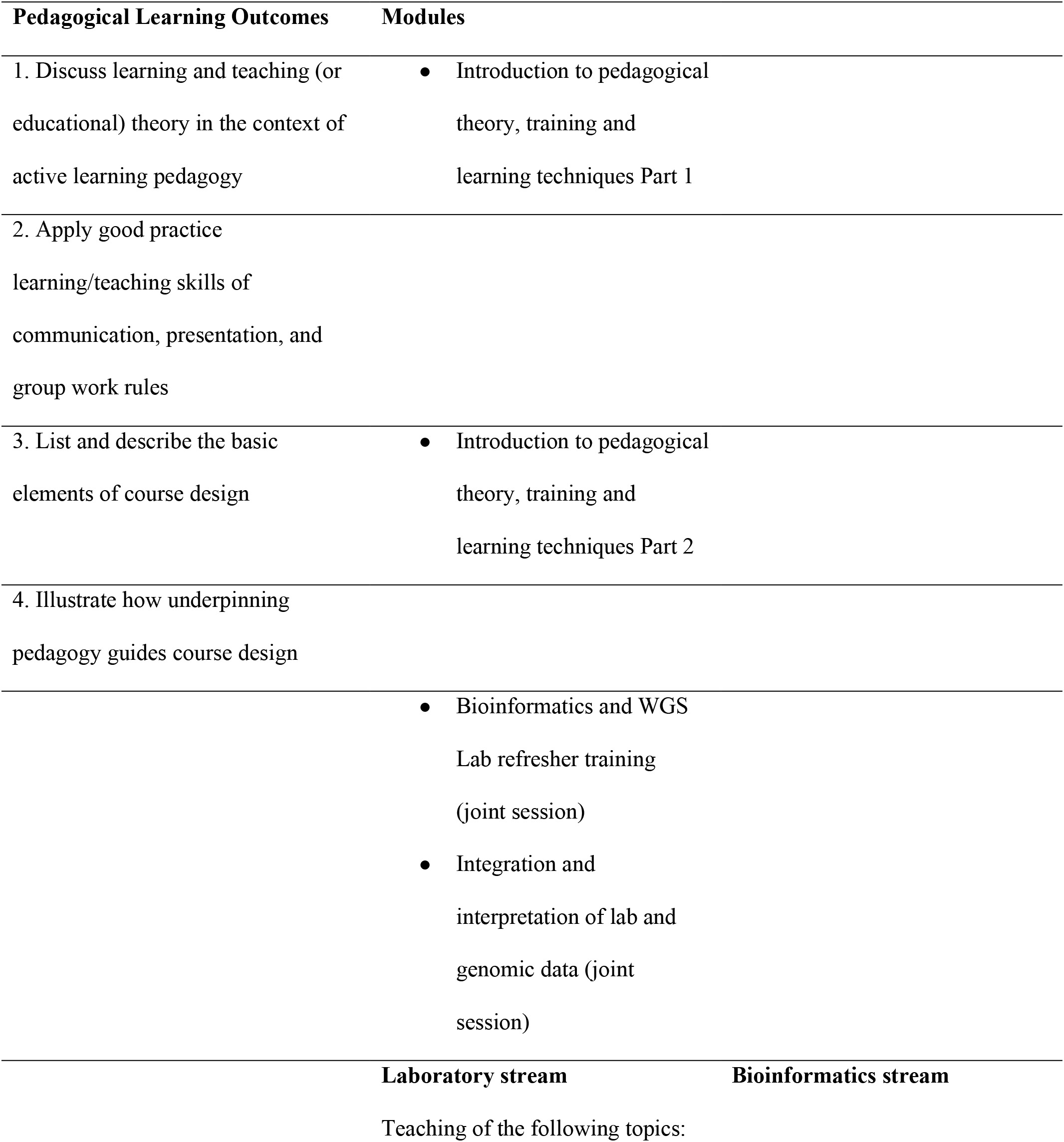
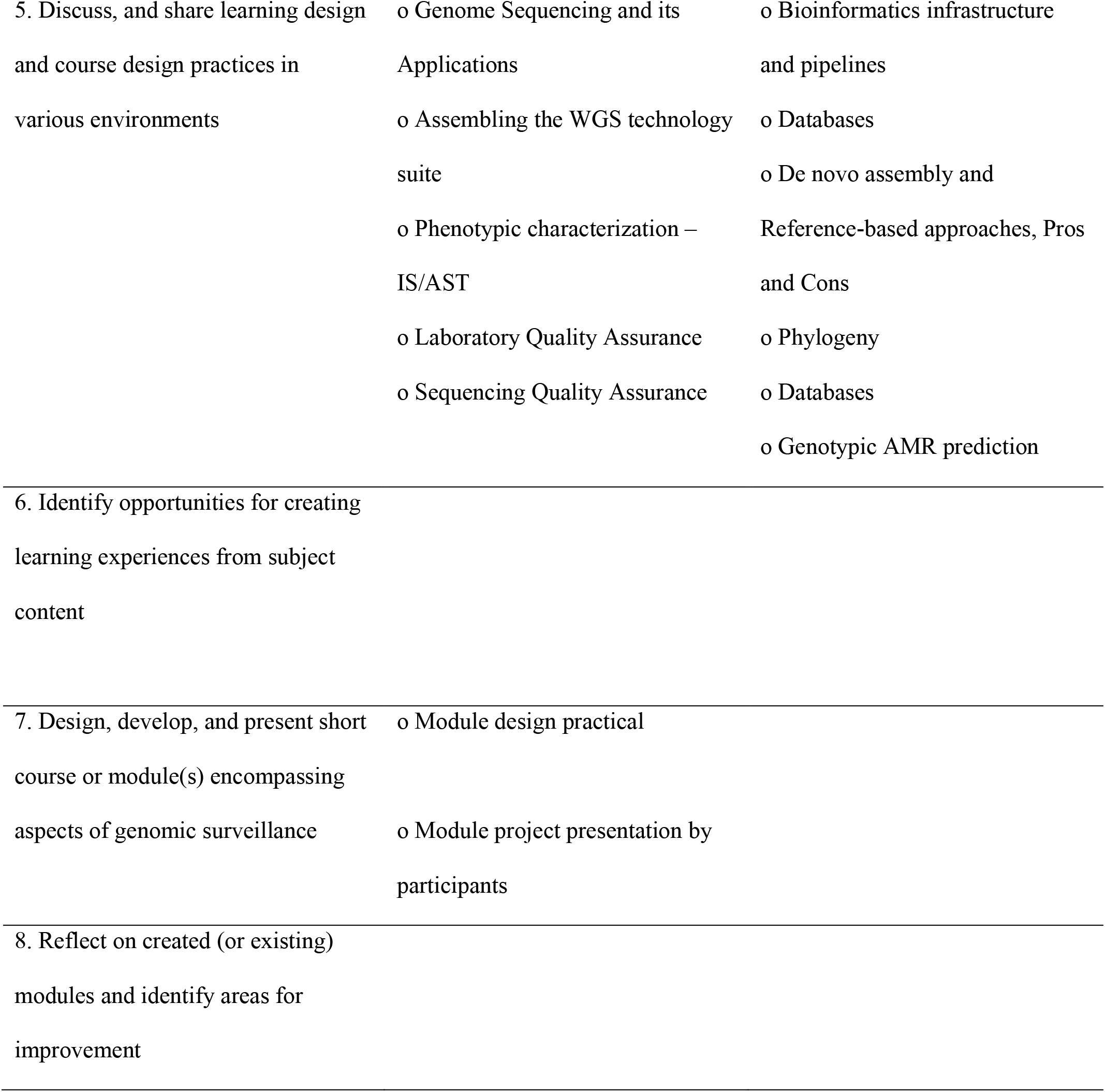
Pedagogical learning outcomes mapped onto content.

### Teaching Approaches

The teaching on the course was integrated and made uniform through the use of templates for instructional design and through the application of the adult and active learning principles throughout [12]. In addition, the instructors used customized teaching approaches to bioinformatics and laboratory topics (Table 2). The main characteristics of bioinformatics and laboratory-specific teaching on this course are Table 2.

**Table 2.** Teaching approaches.

| Strategy | Description | Implementation examples |
| --- | --- | --- |
| **Challenge and problem-based tasks** | Problem-solving and decision-making exercises | • Use card-based blueprint to order processes and procedures in a typical WGS for AMR surveillance pipeline<br>• Use of fictive case scenarios involving training others in genomics aspects |
| **Case studies** | Use of real cases, as published in the literature | • Identify MDR *Salmonella enterica* subsp. *enterica* serotype Kentucky lineage expansion [13]<br>• Investigation of a Brazilian epidemic of *Mycobacterium abscessus* species complex [14]<br>• *Klebsiella pneumoniae* outbreak investigation [15] |
| **Historical and chronological presentation of processes** | Demonstration of lab procedures, visual tours, lab visits, exhibition tour | • Visit to the sequencing facilities at the Sanger Institute<br>• Visit to the exhibition marking the start of the Human Genome Project<br>• Demonstration of library preparation for WGS<br>• Demonstration of best practices for antimicrobial susceptibility testing |
| **Comparative analysis of tools and platforms** | Presenting alternative ways of achieving the same/similar result | • Comparison of data merging and consolidation techniques – interactively with spreadsheets versus programmatically with R and Python<br>• Comparison of different databases and software tools used to determine AMR phenotype based on genomic data |
|  |  | • Comparison of different ways of interpreting bacterial population structures – MLST trees versus phylogenetic trees obtained from WGS data |
| **Project planning and management techniques** | Considering standards and quality control, constraints and limitations, time management; project planning, and constraints (financial, scaling and other) | • Exercise on how to build a computer server that needs to accomplish certain tasks, given a list of hardware components and a fixed budget<br>• Exercise on how to set up a WGS laboratory<br>• QC assessment of libraries and sequencing |
| **Provision of resources** | Provision of links to resources and portals, quality control standards, virtual and cloud tools, toolkits | • Software tools:<br>• AMR prediction tools: ARIBA [16], Pathogenwatch [17], AMRFinder [18]<br>• AMR databases: CARD [19], ResFinder [20]<br>• Nucleotide archives: Genbank [21], European Nucleotide Archive [22]<br>• Data integration tools: dataflo (https://dataflo.io/)<br>• Data visualisation: Microreact [11]<br>• Pipelines: Nextflow [23]<br>• Phylogeny: RAxML [24]<br>• Genome Assembly: SPAdes [25]<br>• QC tools: FastQC [26], Quast [27], Bactinspector [28], Qualifyr [29]<br>• Typing tools: PubMed MLST (www.pubmlst.org)<br>• The Training and Learning Toolkit |
| **Analysis and evaluation** | Consideration and evaluation of participants' specific contexts | • Big-picture exercise where participants evaluate their own circumstances |

### Assessment for Learning

Formative assessment was implemented throughout the course: quizzes, discussions, Q&A sessions, group work and reporting, peer critiquing, module/course presentation with peer and instructors’ feedback, pitching of a genomics plan with instructors’ feedback.

To implement all the learning activities and formative assessment on this course, a number of specific tools, services, and learning material were used, including Post-it Notes, printed cards, bioinformatics online tools and platforms, online quiz tools, presentation tools (PowerPoint, flipcharts), live documents, and lab settings. The classrooms were flexible, with easily movable furniture to accommodate different formats of teaching and learning activities.

### Project-Based Learning

Integrated teaching and learning on this course resulted in the hands-on design of different genomics modules by participants who worked in small groups, so they could collaborate and bring in personal expertise. To guide them with the design, participants used the Toolkit, an online tool shared with the instructors, which provided them with formative feedback during the project and gave them electronic access to course content and outputs throughout the course. The project required the application of the course design principles taught in the pedagogy modules. A well-established methodology was presented, which considers aspects of course design, such as the course aim, target audience, and the level of the course, the formulation of the learning outcomes using revised Bloom’s taxonomy of learning, and outlining of the course content and assessment. How to create specific learning activities taking into account the underpinning pedagogy, and how to plan for the course/module delivery and evaluation, considering specific circumstances, were also discussed. The same process applied in the design, development, delivery, and evaluation of the TtT course was presented to the course participants and used in the final project. This provided the course organizers and instructors with formative feedback. The projects also encouraged creative thinking, collaboration, communication, and presentation skills, and created opportunities for networking.

## FEEDBACK AND EVALUATION

A post-course survey asked participants about course organization, course content, what participants most enjoyed in the course, and any suggestions for changes or improvements. All participants were satisfied with the course materials and indicated that the course had been very relevant to their research or work. Although the course was intensive, participants reported that the modules were well designed and met expectations, and that the combination of active learning strategies was beneficial to their learning and interactions with each other. However, participants also felt that networking and time for discussions could be improved. Since participants were separated by workstream, a suggestion was made to provide a more overlapping structure, whereby bioinformaticians could have an overview of project management approaches for laboratory pipelines. Feedback on the course from organizers, trainers, and trainees was also filmed, and the recording is available online [30].

A follow-up survey was conducted 6 months post-course to understand whether and how participants had applied their knowledge in their research or professional practice. Respondents indicated that they had implemented, or were planning, specific training in microbiology, AMR, sequencing, and bioinformatics techniques to a range of audiences, including university students, epidemiologists, and national-level scientists involved in AMR surveillance. Two of the respondents had already run courses, and more than half have continued to network with other course participants informally. One of the respondents was awarded an international grant to start a pilot training program in their region [31]. Respondents also reported plans or implementation of remodeling their AMR laboratories for WGS and designed SOPs, and tailored WGS data analysis workflows.

### DISCUSSION

This paper describes the first delivery to a regionally diverse audience of a TtT course for building capacity for the application of genomics tools and resources in AMR surveillance. Advances in genomics and emerging infectious disease and environmental threats create a need to train more scientists in pathogen surveillance and bioinformatics skills [32]. This TtT course provided a timely intervention coinciding with global AMR strategies to build capacity in genomic surveillance and the establishment of global centers [33-36].

In LMICs, limited expertise in genomics applications in the areas of epidemiology, surveillance, and public health results in short specialized courses being offered through collaborations with regional and international experts. However, this is severely challenged by the time limitations of trainers and experts, and funding and travel constraints. In addition, public health scientists may not be able to take time away from their core duties or secure the necessary resources to travel for training. This work established a framework to develop skilled local trainers who are knowledgeable about local issues, encouraging local and regional collaboration.

The integrated domain and application-specific approach enabled participants to refresh and reflect on their knowledge and expertise, while learning how to effectively train others. The course applied an innovative combination of teaching approaches, emphasizing the elements of how to teach various audiences in bioinformatics and laboratory techniques used in the genomic surveillance of AMR. While some domain-specific teaching approaches were similar to those more commonly used in general bioinformatics and laboratory education, some of them were uniquely applied to this course (Table 2) [37].

The integration of pedagogy with scientific aspects emphasized that “what to teach” is as important as “how to teach.” Ideally, this should be accompanied by an understanding of how students best learn specific topics, and by reviewing well established methodologies for the training of laboratory techniques or aligning bioinformatics educational approaches with those in computer sciences (e.g. [37]).

The successful implementation of the course was highlighted in the feedback and follow-up evaluation, with participants reporting that they had applied what they learned to establish experimental and analysis pipelines in their institutions and train people in their countries and regions. This highlights the role of the TtT course in building capacity and applying genomics technologies locally with global impact.

Global health partnerships integrate collaborative TtT courses as a key part of effective and sustainable capacity building programs [38]. This TtT course established a scalable model for up-skilling domain expert trainers capable of training others and locally cascading skills development in a context-specific and sustainable way. This provided a foundation to build a global network of trainers and experts, thereby empowering scientists to work more collaboratively on the common goals of tackling AMR.

There is a clear requirement within such courses to establish frameworks for measuring impact and success. To strengthen efforts initiated by participants and other professionals in their local settings, we are working towards building a network of mentors to provide guidance for running courses independently in the future.

## CONCLUSIONS AND FUTURE WORK

Global health consortia and collaborative projects across continents play an important role in establishing centers of excellence, providing infrastructure, facilities, and shared resources for genomic surveillance of AMR [39]. To complement these efforts, training trainers is a sustainable way to build local and regional capacity and networks for research and public health. Our TtT framework presents a potentially effective model for building capacity and developing sustainable trainer networks locally for global impact. To complement this effort, future work will involve the development of standardized trainer resources and best practice guidelines. In addition, digitalization of the course will widen reach across regions where resources and expertise are a major challenge.

## ABBREVIATIONS

AMR: antimicrobial resistance
NIHR: National Institute for Health Research
TtT: Train-the-Trainer
GHRU: NIHR Global Health Research Unit
CGPS: Centre of Genomic Pathogen Surveillance
ACSC: Wellcome Genome Campus Advanced Courses and Scientific Conferences, now Wellcome Connecting Science
WGS: Whole Genome Sequencing
LMIC: low- and middle-income country
LO: learning outcome
SOP: standard operating procedure

## ACKNOWLEDGMENTS

Members of the NIHR Global Health Research Unit for the Genomic Surveillance of Antimicrobial Resistance: Khalil Abudahab, Harry Harste, Dawn Muddyman, Ben Taylor, Nicole Wheeler, and Sophia David of the Centre for Genomic Pathogen Surveillance, Big Data Institute, University of Oxford, Old Road Campus, Oxford, United Kingdom and Wellcome Genome Campus, Hinxton, UK; Pilar Donado-Godoy, Johan Fabian Bernal, Alejandra Arevalo, Maria Fernanda Valencia, and Erik C. D. Osma Castro of the Colombian Integrated Program for Antimicrobial Resistance Surveillance – Coipars, CI Tibaitatá, Corporación Colombiana de Investigación Agropecuaria (AGROSAVIA), Tibaitatá – Mosquera, Cundinamarca, Colombia; K. L. Ravikumar, Geetha Nagaraj, Varun Shamanna, Vandana Govindan, Akshata Prabhu, D. Sravani, M. R. Shincy, Steffimole Rose, and Ravishankar K.N of the Central Research Laboratory, Kempegowda Institute of Medical Sciences, Bengaluru, India; Iruka N Okeke, Anderson O. Oaikhena, Ayorinde O. Afolayan, Jolaade J Ajiboye, and Erkison Ewomazino Odih of the Department of Pharmaceutical Microbiology, Faculty of Pharmacy, University of Ibadan, Oyo State, Nigeria; Celia Carlos, Marietta L. Lagrada, Polle Krystle V. Macaranas, Agnettah M. Olorosa, June M. Gayeta, and Elmer M. Herrera of the Antimicrobial Resistance Surveillance Reference Laboratory, Research Institute for Tropical Medicine, Muntinlupa, the Philippines; Ali Molloy, alimolloy.com; John Stelling, The Brigham and Women’s Hospital; and Carolin Vegvari, Imperial College London.

The authors thank Dr Pamela Black for initiating the program, and the Wellcome Connecting Science Courses team – Yvonne Thornton, Julie Ormond, Martin Aslett – for organizing the course and providing training facilities.

## Figure Legends

These should be on a separate, numbered manuscript sheet. Define all symbols and abbreviations used in the figure. Figures and legends should be intelligible without reading the text of the manuscript.

**Figure 2.**
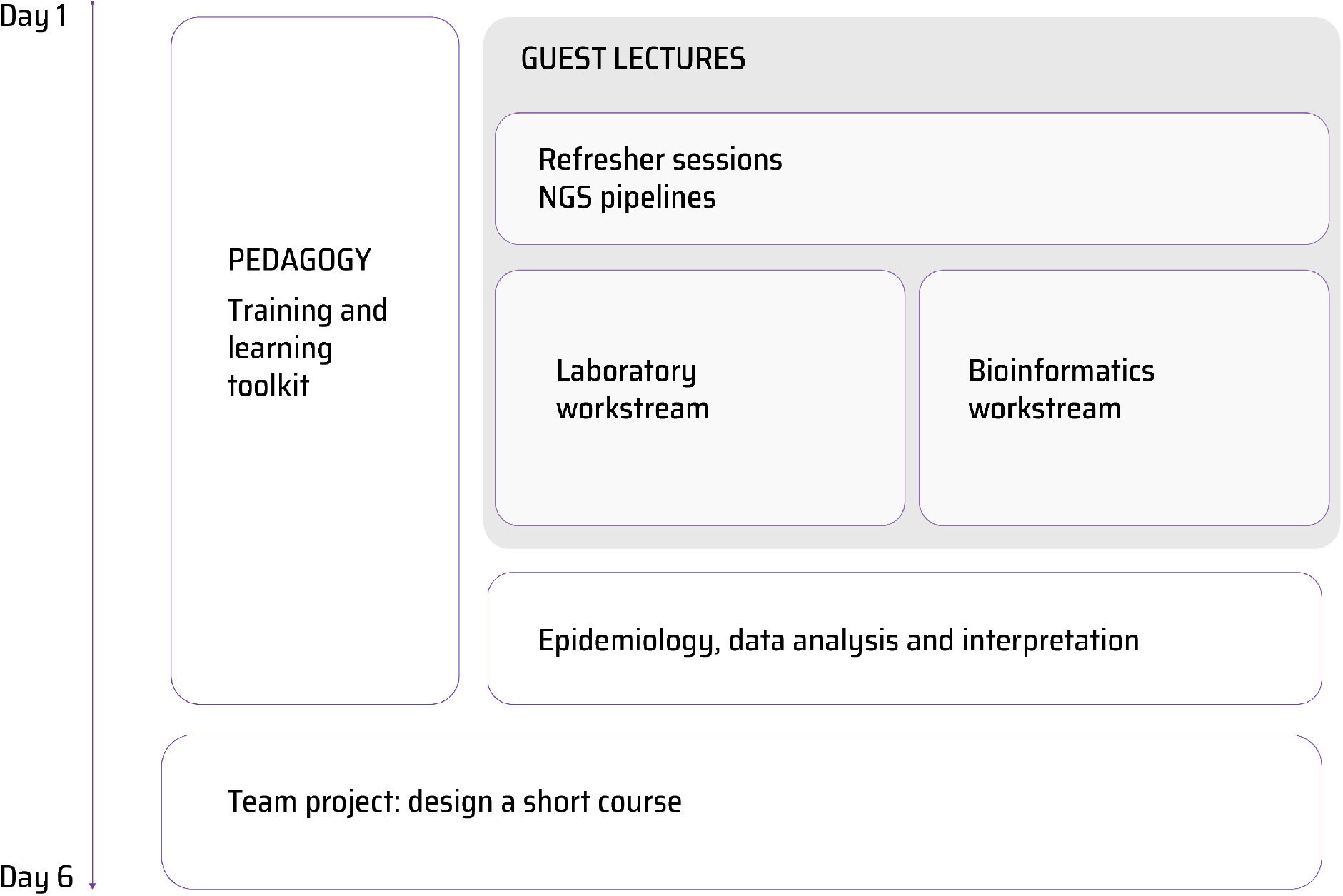
The structure of the course in time, with some modules run in parallel.

